# Optimal structure of heterogeneous stem cell niche: The importance of cell migration in delaying tumorigenesis

**DOI:** 10.1101/082982

**Authors:** Leili Shahriyari, Ali Mahdipour–Shirayeh

**Affiliations:** Mathematical Biosciences Institute, The Ohio State University, OH, USA; Biomedical Research Group, Applied Mathematics Department, University of Waterloo, ON N2L 3G1, Canada

## Abstract

Studying the stem cell niche architecture is a crucial step for investigating the process of oncogenesis and obtaining an effective stem cell therapy for various cancers. Recently, it has been observed that there are two groups of stem cells in the stem cell niche collaborating with each other to maintain tissue homeostasis. One group comprises the border stem cells, which is responsible to control the number of non-stem cells as well as stem cells. The other group, central stem cells, regulates the stem cell niche. In the present study, we develop a bi-compartmental stochastic model for the stem cell niche to study the spread of mutants within the niche. The analytic calculations and numeric simulations, which are in perfect agreement, reveal that in order to delay the spread of mutants in the stem cell niche, a small but non-zero number of stem cell proliferations must occur in the central stem cell compartment. Moreover, the migration of border stem cells to the central stem cell compartment delays the spread of mutants. Furthermore, the fixation probability of mutants in the stem cell niche is independent of types of stem cell division as long as all stem cells do not divide fully asymmetrically. Additionally, the progeny of central stem cells have a much higher chance than the progeny of border stem cells to take over the entire niche.

## Introduction

Many studies have been designed to investigate the stem cell dynamics because of their crucial role in many diseases such as cancer. Recently, there have been several studies about employing stem cell therapy for various diseases including cancers [1]. In an in vivo study, it has been observed that the injection of rat umbilical cord stem cells (rUSCs) can completely abolish rat mammary carcinomas with no evidence of metastasis or recurrence hundred days post-tumor cell inoculation [2].

Knowing stem cell dynamics such as their division and death rates, their types of divisions, and the rate of each type of division can assist us to improve stem cell therapies. This knowledge can suggest ways of altering the structure of the stem cell niche to minimize the number of mutant stem cells and control the cells’ growth rate. Understanding the division patterns of stem cells in healthy and malignant tissues can also help us to determine the origins of several diseases like cancer.

Tissue cells are divided in two general categories: Stem cells and non-stem cells, which include transit amplifying (TA) and fully differentiated (FD) cells. The main characteristic of stem cells is their diverse division types. There are two types of stem cell divisions: symmetric and asymmetric. When a stem cell divides asymmetrically, one of the newborn cells is a stem cell and the other one is a non-stem cell. There are two types of stem cell symmetric division: proliferation (both newborn cells are stem cells) and differentiation (both children are TA cells). It has been shown that normal adult human stem cells divide mostly symmetrically in the intestinal crypts [3]. There is also evidence showing that stem cells divide asymmetrically in the response to the injury during tissue regeneration [4]. Since the asymmetric stem cell division leads to a high probability of two-hit mutant productions [5], injuries might increase the chance of tumorigenesis.

There are many experimental observations about the cooperation among different cell types within and around the niche. For instance, in the intestinal crypts, Lgr5 stem cells receive niche support from Paneth cells, which are located at the bottom of the crypts in the stem cell niche (SC niche) [6]. The cooperation of epithelial hair follicle stem cells (HFSCs) and melanocyte stem cells (MCSCs) in the bulge niche leads to the differentiation and proliferation of MCSCs [7]. There is evidence showing that the haematopoietic stem cell (HSC) niche in the bone marrow is protected and regulated by the HSC progeny and mesenchymal stem cells [8–10]. Additionally, two distinct groups of dormant (d-HSCs) and activated (a-HSCs) stem cells have been identified [11–14].

It has been suggested that stem cells in many tissues are characterized by a bi-compartmental organization [15]. One of the compartments is responsible to regenerate the tissue, while the other compartment controls the SC niche. Such patterns have been observed in several adult SC niches, such as those in hair follicles, blood, intestine, and brain [15]. More details of the stem cell dynamics in the bi-compartmental niche structure have been recently reported by Ristma et al. [16]. They observed that there are two groups of stem cells in the SC niche: border stem cells (BSCs) and central stem cells (CeSCs). Border stem cells, which are located between central stem cells and transit amplifying cells, mostly differentiate to control the number of non-stem cells. The central stem cells, which are located at the middle of stem cell niche surrounded by BSCs, mostly proliferate to control the number of stem cells [16].

Mathematical models have been developed to understand cell dynamics in both normal an malignant tissue [17–21]. Many of these models investigate the dynamics of stem cells [22–28]. Some of the studies are concentrated on mutations in stem cells, because of their importance in the initiation and progressions of cancers, see e.g. [29–34]. Shahriyari and Komarova [35] developed a stochastic bi-compartmental model for the SC niche to obtain the optimal structure of the SC niche, which delays two-hit mutants’ production. They found that the optimal structure corresponds to the stem cell symmetric divisions and most stem cell divisions occurring in BSCs (very small but not zero number of division happening in the CeSCs). Recently, we have developed a 4-compartmental stochastic model, which includes two stem cell groups, to study the movement of mutants in the intestinal and colon crypts [36].

Ritsma et al. [16] observed a small number of migrations from BSCs to CeSCs. However, the model developed in [35] does not consider the possibility of the migration from BSCs towards the CeSC compartment. In this work, by generalizing that model, adding the possibility of migration from BSCs to CeSCs, we obtain the optimal niche’s architecture that delays the spread of mutants in the niche. Our model reveals that the migration from BSCs to CeSCs delays the spread of mutants. The model also shows that the percentage of symmetric and asymmetric divisions does not make a difference in the probability of mutants taking over the entire SC niche as long as all stem cell divisions are not fully asymmetric. The number of mutants does not change if all stem cell divisions are asymmetric, because when a mutant stem cell divides asymmetrically it renews itself and it produces a mutant non-stem cell. Furthermore, the model indicates that it is unlikely that the progeny of mutant BSCs take over the entire niche.

## Results

### Model set-up

We develop a Moran model for cell dynamics in the SC niche by generalizing the model introduced in [35]. The model contains two stem cell compartments: the BSC compartment (*S*_*b*_) and the CeSC compartment (*S*_*c*_). In this model, we only keep track of stem cells and denote the population size of CeSCs and BSCs respectively by |*S*_*c*_| and |*S*_*b*_|.

In the Moran cell dynamics models, at each updating time step one cell dies and one cell divides to keep the number of cells constant. Here, we assume two non-stem cells die and they are replaced by two stem cells. The missing non-stem cells could be replaced by the divisions of non-stem cells. Since we are modeling the stem cell dynamics, we only consider those that are replaced by stem cell divisions. In other words, we assume at each updating time step, two non-stem cells are required to be produced by some stem cell divisions. We choose two cell divisions only to accommodate stem cells’ symmetric divisions and keep the number of cells at each compartment canstant, because we are constructing a Moran model to obtain fixation probabilities. In this model, at each updating time step two cells are randomly chosen to divide based on their fitness. We assume that the normalized fitness of wild-type cells is 1, while the relative fitness of mutants is *r*. We consider a range of values for the mutants’ fitness: *r* < 1 (disadvantageous mutants), *r* > 1 (advantageous mutants), and *r* = 1 (neutral mutants). For instance, if a division occurs in the BSC compartment, with probability 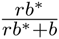 one BSC mutant divides, and with probability 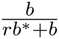 one normal cell divides, where *b* and *b\** are respectively the number of normal and mutant BSCs. At each updating time, two stem cell divisions happen based on the following algorithm (See Fig 1).

- With probability 1 – *σ*, two BSCs divide asymmetrically.
- Or, with probability *σ*, one BSC differentiates to produce two non-stem cells.

- And, with probability *γ*, one CeSC proliferates. Then, one random cell from CeSc migrates to the *S*_*b*_ compartment, to replace the differentiated BSC.
- Or, with probability 1 – *γ*, one border stem cell proliferates to replace the differentiated BSC cell.

*And, with probability *α,* one random cell from the *S*_*b*_ compartment migrates to the *S*_*c*_ compartment. Then, one random cell from the *S*_*c*_ group migrates to the *S*_*b*_ compartment, to keep the number of cells at each compartment constant.

**Fig. 1.**
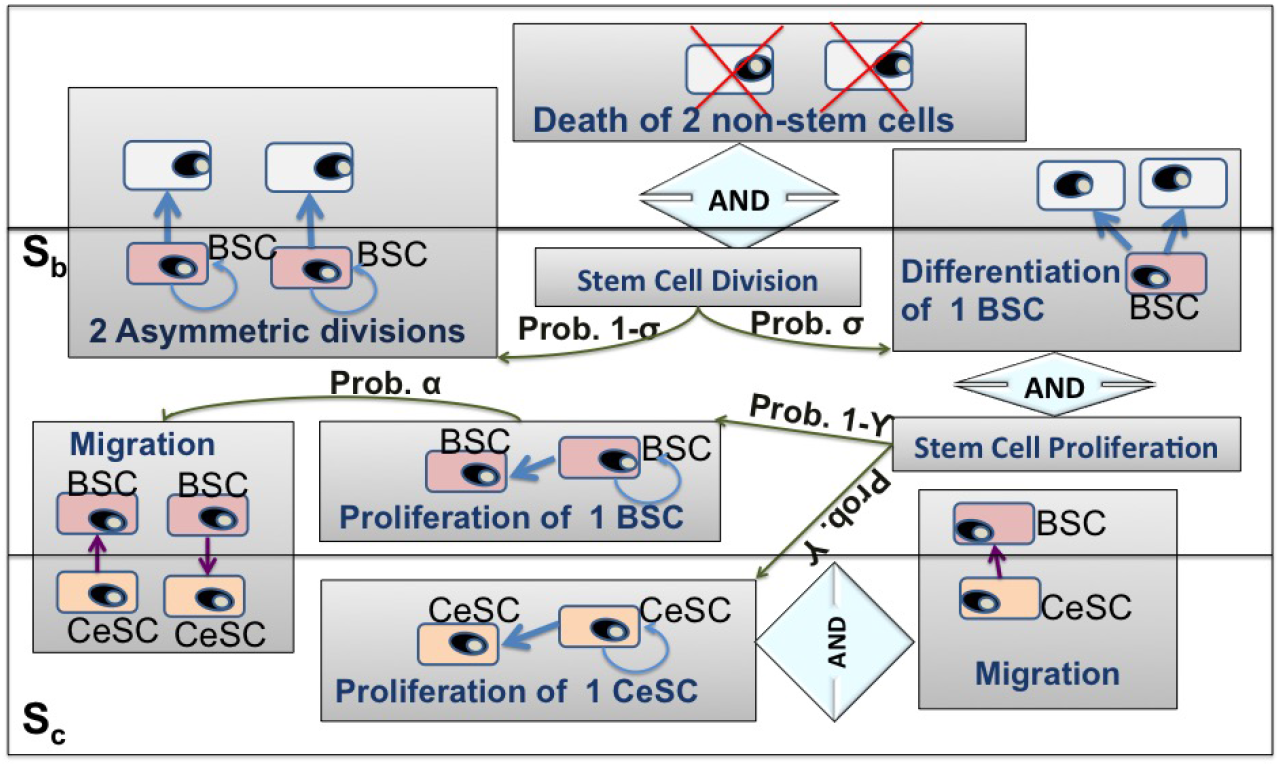
Schematic representation of the model. When two non-stem cells die, either two BSCs divide asymmetrically or one BSC differentiate to generate two non-stem cells. When a BSC cell differentiates, a proliferation occurs in the SC niche to replace the differentiated BSC cell.

The parameters introduced in the above algorithm are given in Table 1. Also, the general mechanism, which is illustrated in the algorithm, can be seen in Figure 1, in which the different steps of the procedure are given in detail.

**Table. 1.** Model parameters.

| Symbol | Description |
| --- | --- |
| $r$ | Prolif. rate of malignant w.t. and stem cells |
| $\sigma$ | Prob. of symmetric division in CeSCs |
| $\gamma$ | Prob. of proliferation in CeSC |
| $\alpha$ | Prob. of migration from BSC to CeSc when a BSC proliferates |
| $\vert S_c\vert $ | Total number of CeSCs |
| $\vert S_b\vert $ | Total number of BSCs |

### The progeny of mutant CeSCs will take over the CeSC compartment and the entire SC niche with a very high probability

The numerical simulations and analytic calculations illustrated in the Materials and Methods section show that the fixation probability of CeSC mutants in the CeSC compartment is an increasing function of the mutants’ fitness *r.* Furthermore, the probability that the progeny of CeSC advantageous mutants will take over the *S*_*c*_ compartment is very high. Furthermore the analytic results and simulations show that this probability is not very sensitive to *α*, when *γ* (the probability that CeSCs proliferate) is large or when the mutants are not advantageous (Figure 2).

**Fig. 2.**
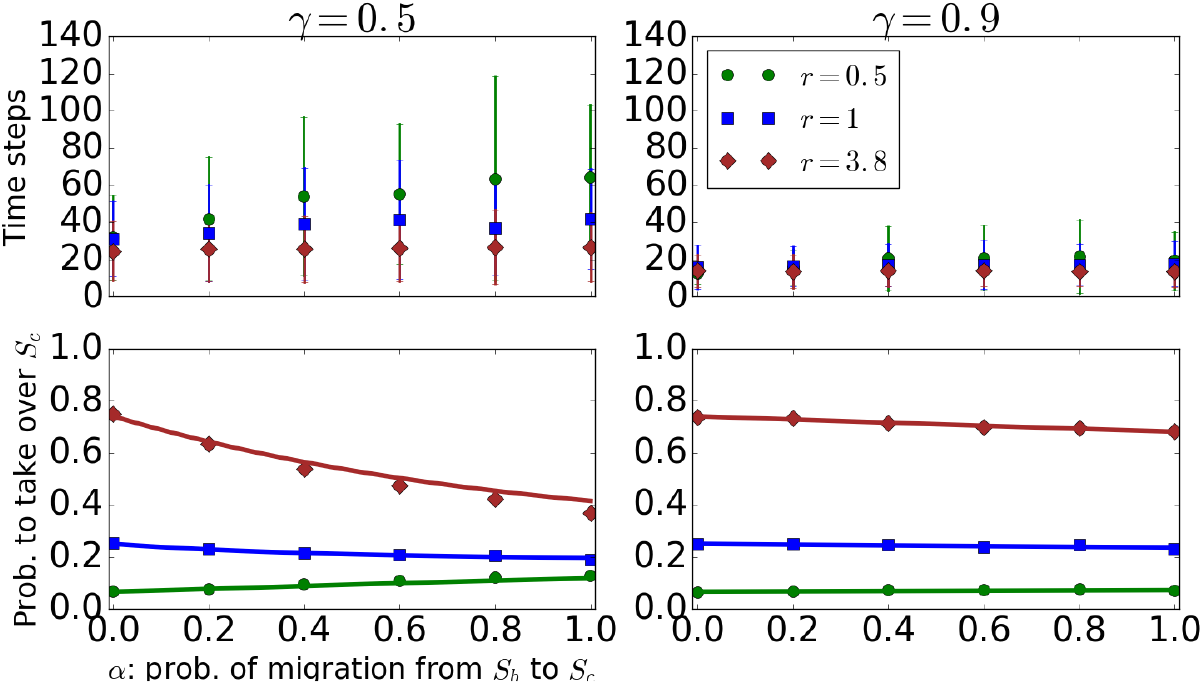
The probability that the progeny of a CeSC mutant taking over the CeSC compartment. The bottom sub-figures show the mean and standard deviations of results of simulations (points and bars) and results of formula (7) (solid lines) (solid lines) for the probability that the progeny of a CeSC mutant will take over the CeSCs. The top sub-figures represent the results of simulations for the time that the progeny of one mutant CeSC needs to take over the *S*_*c*_ compartment. The parameters for this figure are *σ* = 1, |*S*_*c*_| = 4, and |*S*_*b*_| = 7.

The results of simulations, which are in perfect agreement with analytic calculations (Figures 2 and 3), show that with a high probability the progeny of CeSC mutants will take over the entire SC niche. Moreover, this probability is independent of *σ* (the probability of symmetric division). The time to fixation of advantageous mutants increases and their fixation probability decreases, when the migration rate from *S*_*b*_ to *S*_*c*_ rises (see Figure 2).

**Fig. 3.**
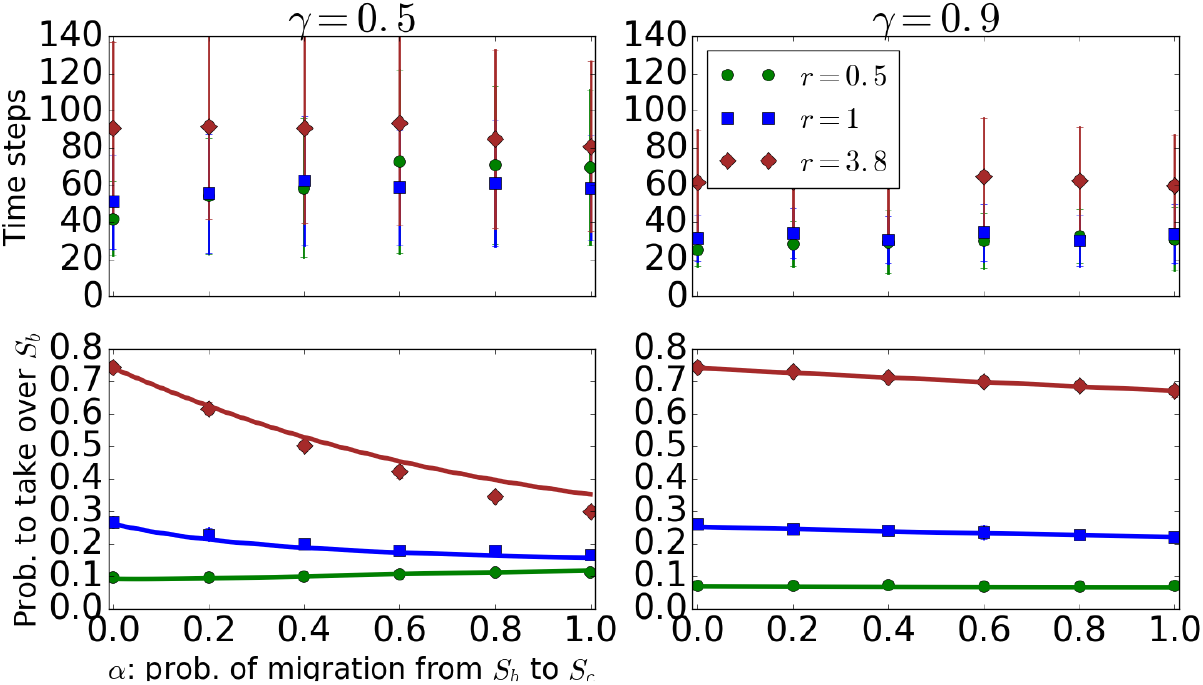
The probability that the progeny of a CeSC mutant taking over the BSC compartment. The bottom sub-figures represent the mean and standard deviations of results of simulations (points and bars) and results of formula (10) (solid lines) for the probability that the progeny of a mutant CeSC will take over BSCs. The top sub-figures show the results of simulations for the time that the progeny of one mutant CeSC needs to take over the *S*_*c*_ compartment. The parameters are *σ* = 1, |*S*_*c*_| = 4, and |*S*_*b*_| = 7.

**Fig. 4.**
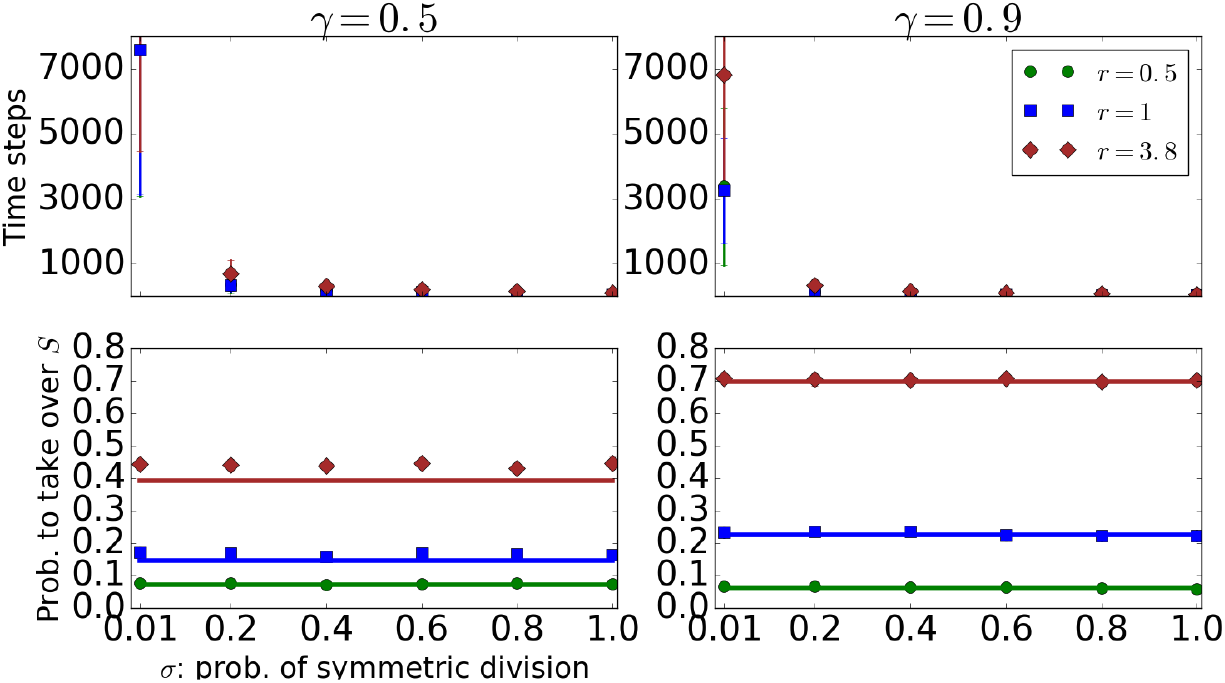
The time and probability that the progeny of a CeSC mutant taking over the SC niche. The bottom sub-figures represent the mean and standard deviations of results of simulations (points and bars) and the resuts of formula (2) (solid curves) for the probability that the progeny of a mutant CeSC will take over the SC niche as a function of *σ*, the probability of symmetric division. The top sub-figures show the results of simulations for the time that the progeny of one mutant CeSC needs to take over the niche. The parameters are *α* = 0.5, |*S*_*c*_| = 4, and |*S*_*b*_| = 7.

### It is unlikely that the progeny of a BSC mutant take over the BSC compartment and the entire SC niche

The analytic calculations and simulations, which are in perfect agreement, show that the fixation probability of BSC mutants in BSC and CeSC compartments is an increasing function of *α,* the probability of migration from *S*_*b*_ to *S*_*c*_ (Figures 5 and 6). When *α* = 0, BSCs do not have a chance to move to the CeSC compartment. Therefore, with a high chance BSC mutants will differentiate and be removed from the niche. Furthermore, advantageous mutants differentiate fast, because with a high probability they will be chosen to divide (i.e. differentiate). Since differentiations only happen in the BSC compartment, the probability that the progeny of a BSC mutant will take over the entire SC niche is very small. For example, the fixation probability of one BSC mutant in the BSC and CeSC compartments are approximately zero when *α* = 0, and these probabilities are around 0.05 when *α* = 1 for *γ* = 0.5 (Figures 5, 6, and 7). Although the time that BSC mutants need to take over the niche is increasing when stem cells divide mostly asymmetrically. The fixation probability of BSC mutants in the entire SC niche does not depend on *σ,* the probability of symmetric division, when *σ* > 0 (Figure 7).

**Fig. 5.**
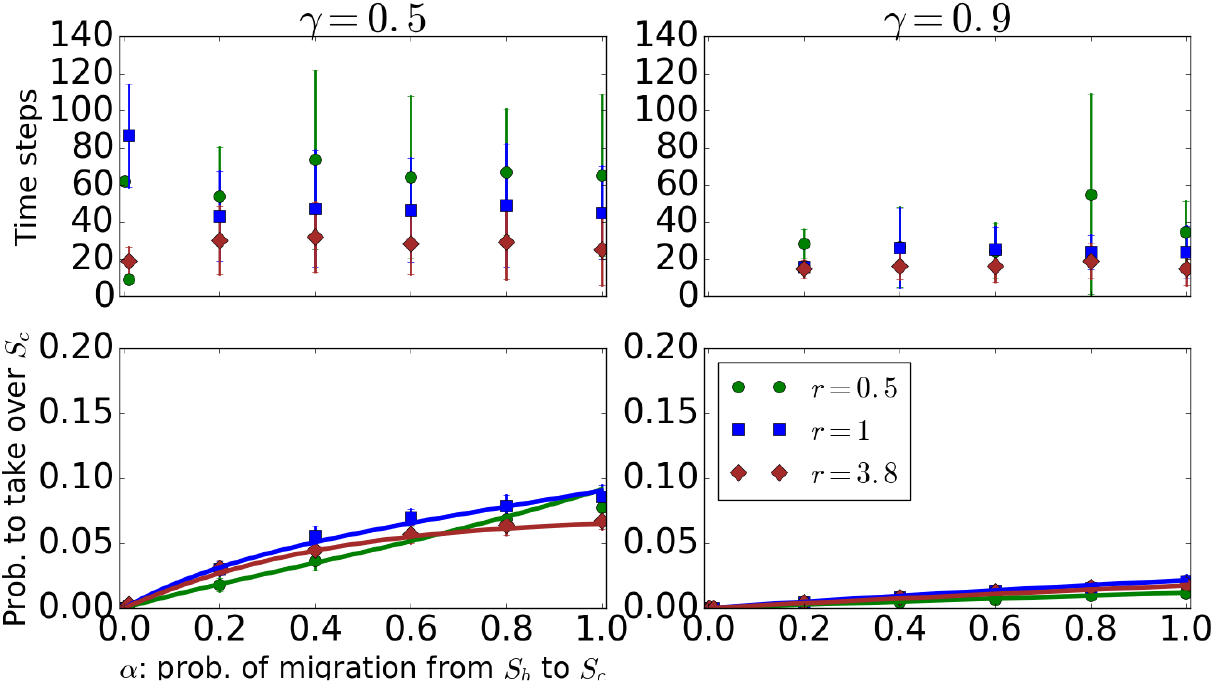
The probability that the progeny of a BSC mutant taking over the CeSC compartment. The bottom sub-figures indicate the mean and standard deviations of results of simulations (points and bars) and results of formula (7) (solid lines) for the probability that the progeny of a mutant BSC will take over the CeSCs. The top sub-figures represent the results of simulations for the time that the progeny of one mutant BSC needs to take over the *S*_*c*_ compartment. The parameters for this figure are *σ* = 1, |*S*_*c*_| = 4, and |*S*_*b*_| = 7.

**Fig. 6.**
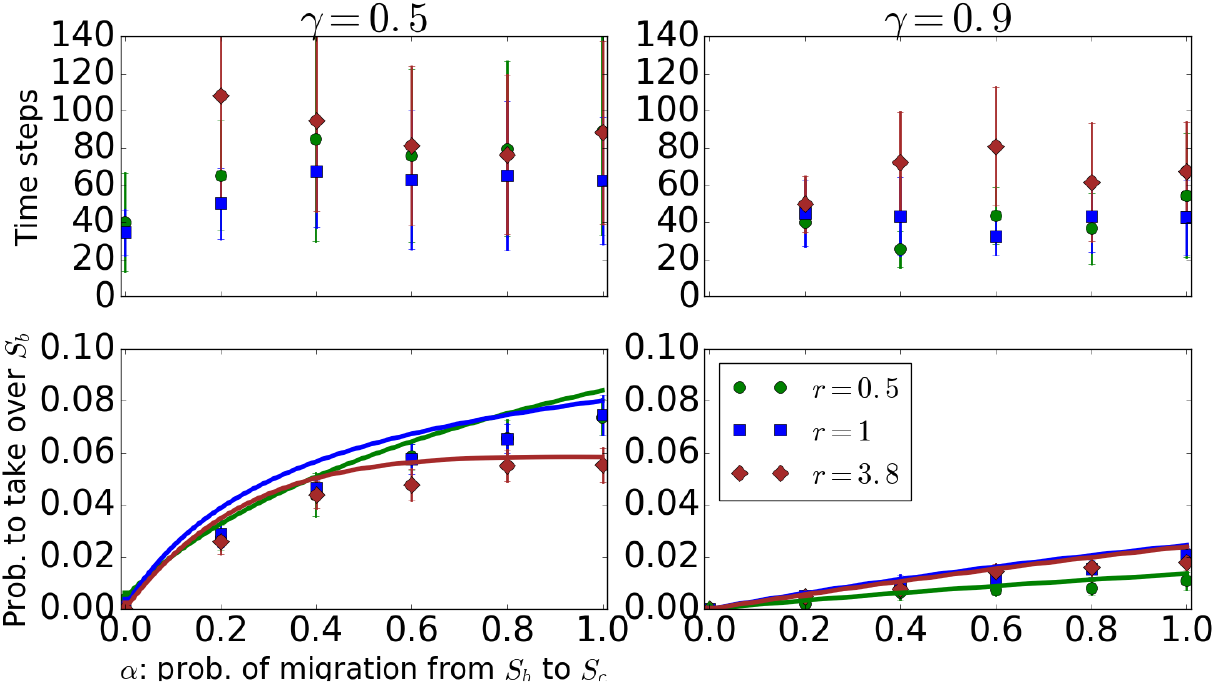
The time and probability that the progeny of a BSC mutant taking over the BSC compartment. The bottom sub-figures represent the mean and standard deviations of results of simulations (points and bars) and results of formula (10) (solid lines) for the probability that the progeny of a mutant BSC will take over the BSCs. The top sub-figures show the results of simulations for the time that the progeny of one mutant BSC needs to take over the *S*_*b*_ compartment. The parameters for this figure are *σ* = 1, |*S*_*c*_| = 4, and |*S*_*b*_| = 7.

**Fig. 7.**
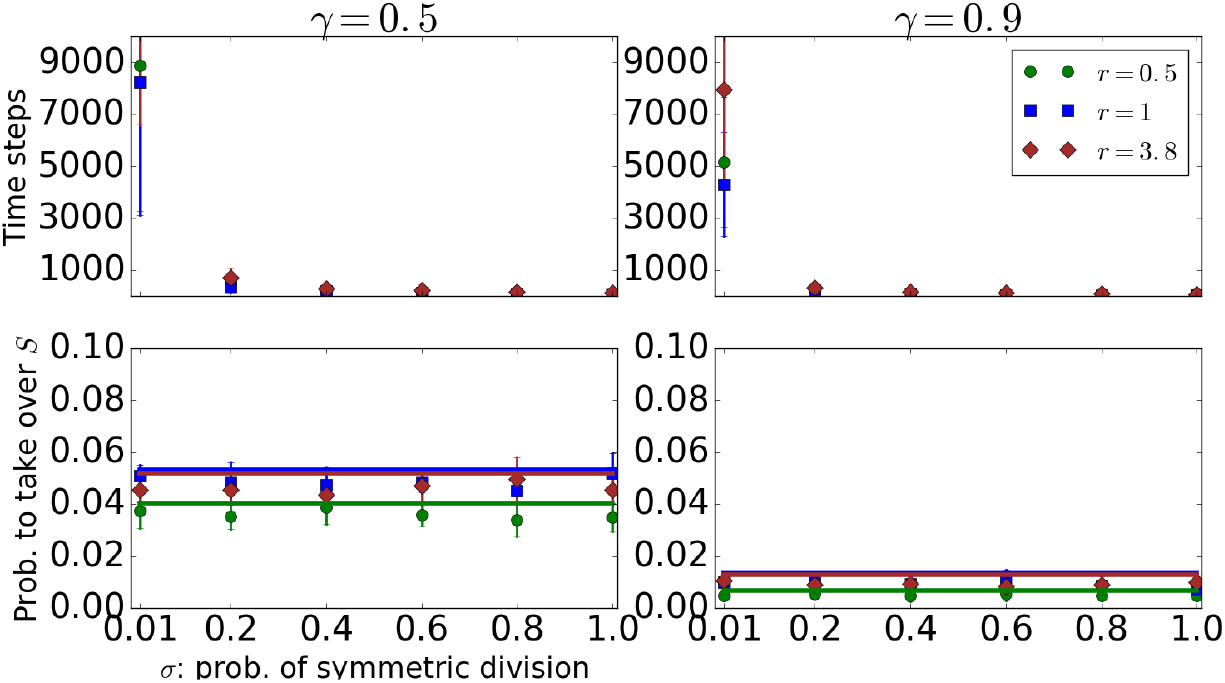
The time and probability that the progeny of a BSC mutant taking over the SC niche. The bottom sub-figures represent the mean and standard deviations of results of simulations (points and bars) and results of formula (2) (solid lines) for the probability that the progeny of a mutant BSC will take over the SC niche as a function of *σ*, the probability of symmetric division. The top sub-figures show the results of simulations for the time that the progeny of one mutant BSC needs to take over the niche. The parameters are *α* = 0.5, |*S*_*c*_| = 4, and |*S*_*b*_| = 7.

### The fixation probability of mutants does not depend on the probability of migration from *S*_*b*_ to *S*_*c*_ when *γ* is close to one

Simulations and analytic calculations show that the fixation probability of mutants in the SC niche is an increasing function of *α,* the probability of migration from *S*_*b*_ to *S*_*c*_, when *γ* is not large. However, the fixation probability of mutants in the niche is not sensitive to *α* when most of the stem cell proliferations occur in the CeSC compartment, i.e. *γ* > 0.9. BSCs have a chance to migrate to the CeSC compartment when proliferations occur in the BSC compartment. Thus, when most of the proliferations happen in the CeSC compartment, then BSCs have a very small chance to migrate from *S*_*b*_ to the CeSC compartment. This leads to the probability of the progeny of a mutant stem cell taking over the niche becoming almost insensitive to *α* when *γ* is large (Figures 2, 3, 5, and 6).

### Migration from *S*_*b*_ to *S*_*c*_ delays tumorigenesis

The fixation probability of neutral and disadvantages CeSC mutants in the entire SC niche is not very sensitive to *α,* which is the probability of migration from *S*_*b*_ to *S*_*c*_. However, the fixation probability of advantageous CeSC mutants is decreasing function of *α.* Moreover, simulations show that the time to fixation of CeSC mutants in the SC niche is an increasing function of *α* for advantageous mutants (Fig 8). In other words, a higher rate of migration from *S*_*b*_ to *S*_*c*_ maximizes the time that the progeny of advantageous CeSC mutants need to take over the SC niche. Therefore, the optimal value for *α* in terms of delaying the spread of advantageous CeSC mutants is one.

**Fig. 8.**
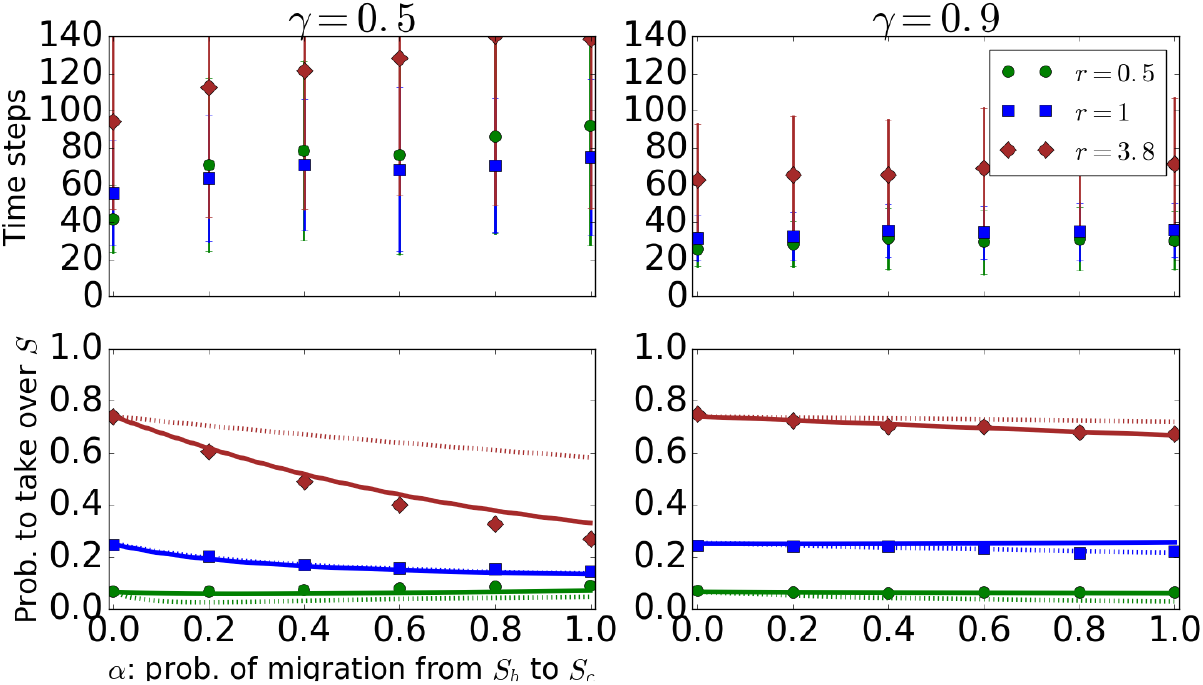
The time and probability that the progeny of a CeSC mutant taking over the entire SC niche. The bottom sub-figures represent the mean and standard deviations of results of simulations (points and bars), results of formula (26) (solid lines), and approximated formula (12) (dashed lines) for the probability that the progeny of a mutant CeSC will take over the niche. The top sub-figures show the results of simulations for the time that the progeny of one mutant CeSC needs to take over the entire niche. The parameters for this figure are *σ* = 1, |*S*_*c*_| = 4, and |*S*_*b*_| = 7.

**Fig. 9.**
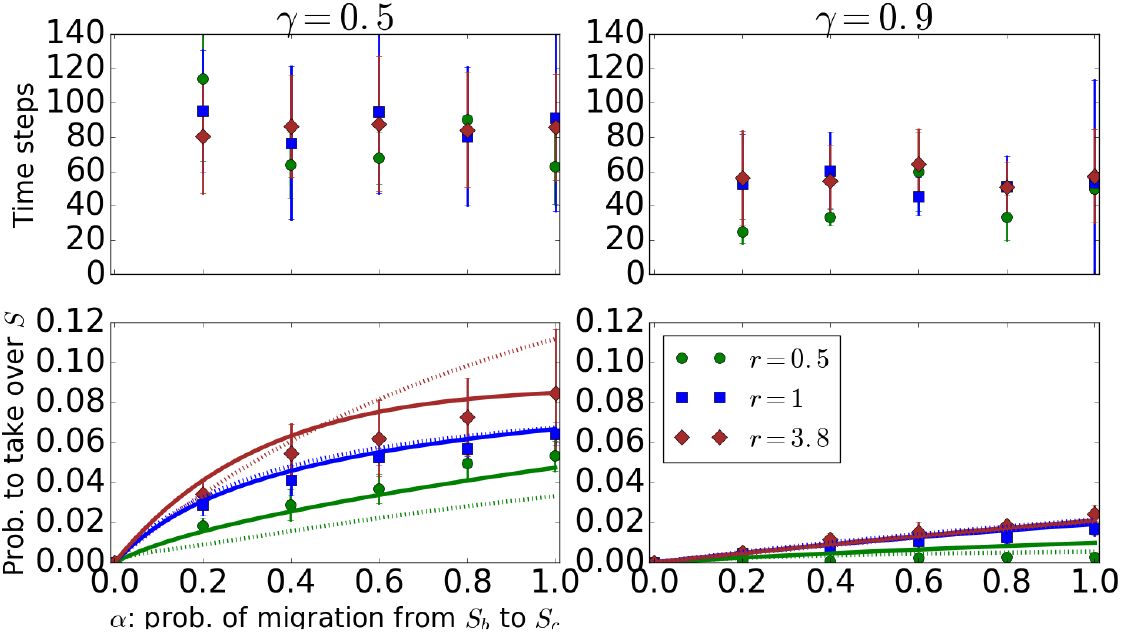
The time and probability that the progeny of a BSC mutant taking over the entire SC niche. The bottom sub-figures represent the mean and standard deviations of results of simulations (points and bars), results of formula (26) (solid lines), and approximated formula (12) (dashed lines) for the probability that the progeny of a mutant BSC will take over the niche. The top sub-figures show the results of simulations for the time that the progeny of a mutant BSC needs to take over the entire niche. The parameters for this figure are *σ* = 1, |*S*_*c*_| = 4, and |*S*_*b*_| = 7.

### The fixation probability of mutants in the SC niche is independent of the probability of symmetric division (*σ*) when *σ* > 0

Although the symmetric division of stem cells minimizes the probability of double-hit mutant production [5, 37], the fixation probability of mutants in the SC niche is independent of the rate of symmetric stem cell division (*σ*) as long as stem cells do not divide fully asymmetrically (*σ ≠* 0) (Figures 4 and 7). When no symmetric proliferation occurs in the SC niche, then the mutant stem cells do not have a chance to spread in the niche. Furthermore, the time that mutant stem cells need to take over the stem cell niche is a decreasing function of the probability of symmetric division *σ*. In other words, asymmetric division delays the process of spreading mutants in the stem cell niche.

### Optimal structure

Figure 10 (a)–(c) represents a contour plot for the probability of a CeSC mutant taking over the SC niche varying *α,* which is the probability of migration from *S*_*b*_ to *S*_*c*_, and *γ*, which is the probability of proliferation in *S*_*c*_. The fixation probability of CeSC mutants is zero when *γ* = 0, because in this case no division will happen in the CeSC compartment. When *γ* ≈ 0, the minimum fixation probability of a disadvantageous CeSC mutant in the entire SC niche corresponds to *α* ≈ 0. However, if *γ* ≠ 0, then the probability that the progeny of an advantageous or neutral CeSC mutant will take over the entire niche is minimized when *α* > 0.2 and *γ* ≠ 0. Additionally, when *γ* ≫ 0, the minimum value of the fixation probability of CeSc mutants in the entire SC niche corresponds to the *α* = 1. Note, when *α* = 1, BSCs have a high chance to migrate to CeSCs, therefore CeSC mutants gain a high chance to move to *S*_*b*_, because each migration from *S*_*b*_ to *S*_*c*_ is coupled with a migration from *S*_*c*_ to *S*_*b*_. Then, with a high probability, mutants in the *S*_*b*_ compartment will differentiate and will be removed from the SC niche.

**Fig. 10.**
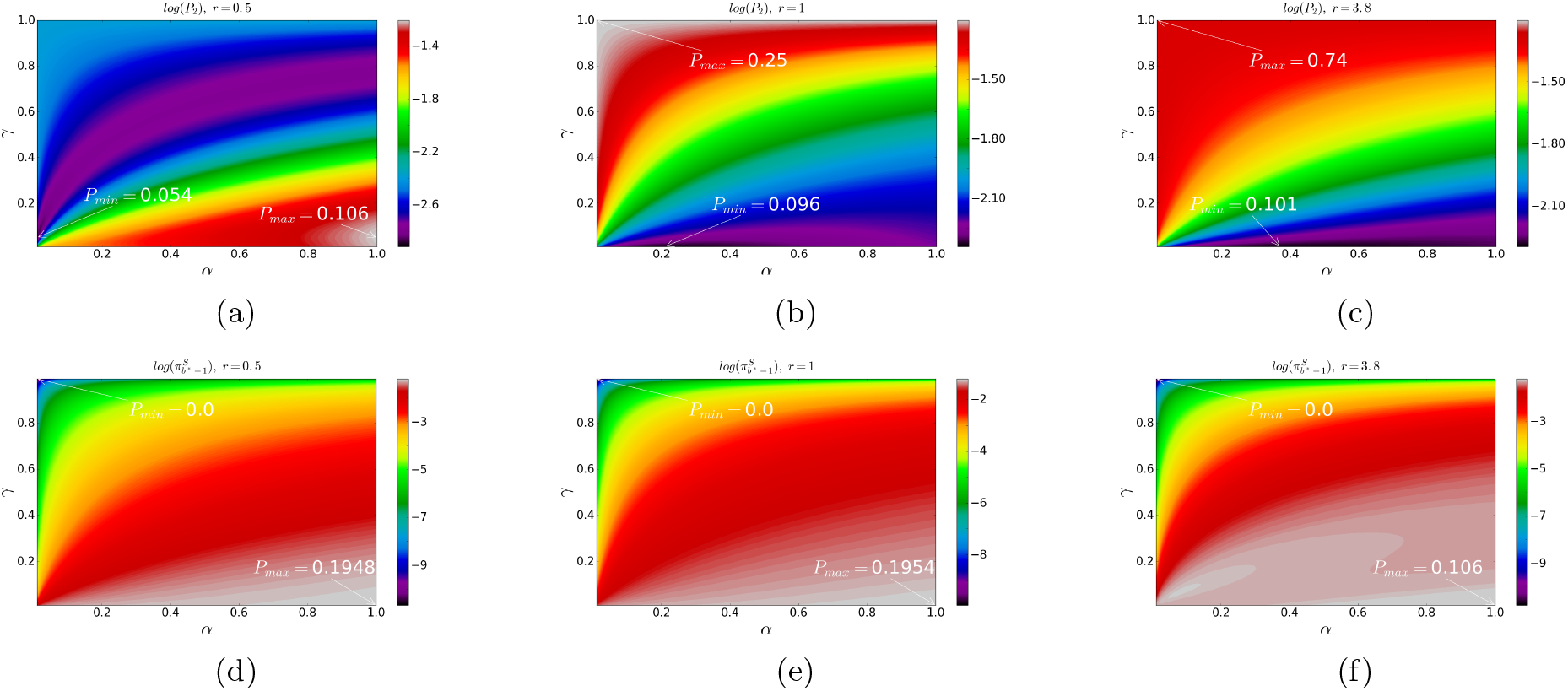
Contour plot of the fixation probability of an initial mutant in the entire SC niche. Sub-figures (a)-(c) show the probability of the progeny of a *S*_*c*_ mutant in taking over the entire SC niche. Plots (d)-(f) indicate the probability of the progeny of a *S*_*b*_ mutant taking over the entire SC niche. Parameters are |*S*_*c*_| = 4, |*S*_*b*_| = 7, (a,d): *r* = 0.5, (b,e): *r* = 1, and (c,f): *r* = 3.8.

**Fig. 11.**
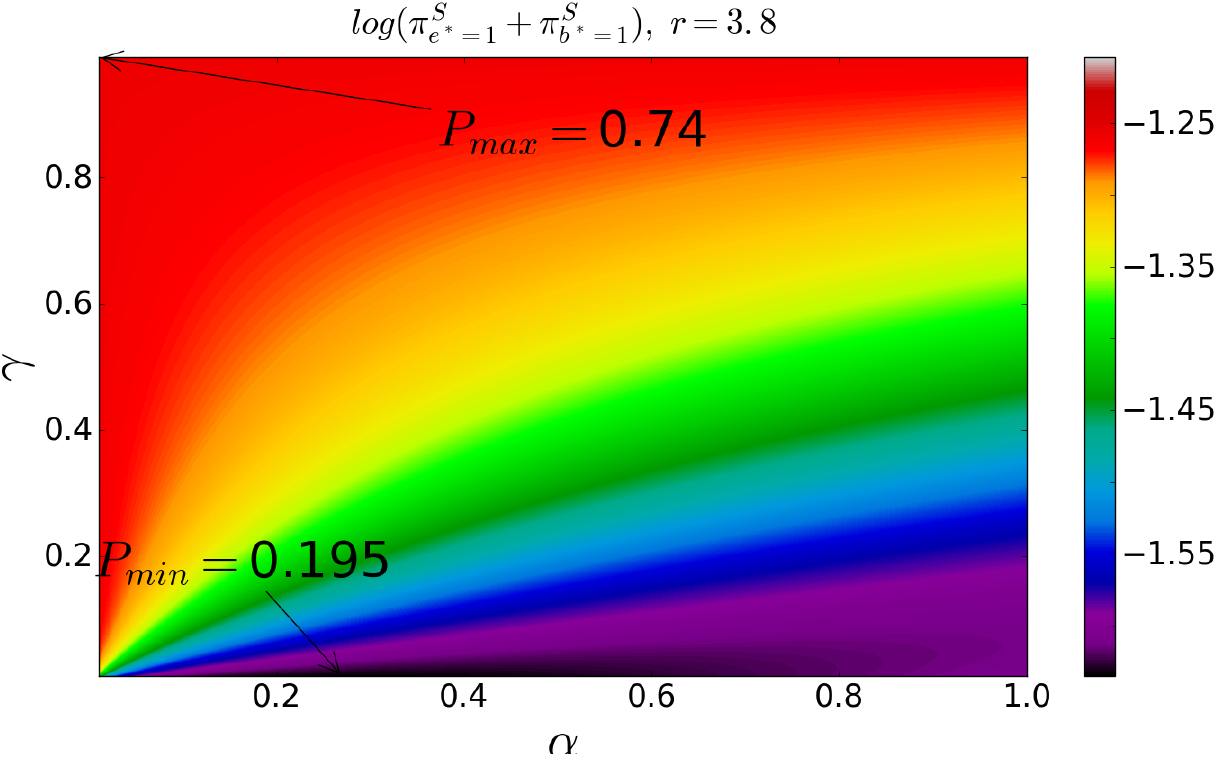
Contour plot of the summation of fixation probability of a CeSC mutant and fixation probability of a BSC mutant in the entire SC niche. This figure shows the summation of the probability of the progeny of a *S*_*c*_ mutant taking over the entire SC niche and the probability of the progeny of a *S*_*b*_ mutant taking over the entire SC niche. Parameters are |*S*_*c*_| = 4, |*S*_*b*_| = 7, and *r* = 3.8.

Figure 10(d)–(f) indicates the probability of a BSC mutant taking over the entire SC compartment, which is very small, as a function of *α* and *γ*. In this case the minimum fixation probability, which is zero, happens when *γ* = 1, because in this case, all stem cell proliferations occur in the CeSC compartment and BSCs only differentiate. When *γ* < 1, the fixation probability of BSC mutants in the entire SC niche is an increasing function of *α.* The maximum of the probability that the progeny of a mutant BSC cell will take over the entire SC, which is around 0.2 for neutral and disadvantageous mutants and around 0.1 for advantageous mutants, happens when *γ* = 0 and *α* = 1.

From the figure 11, which shows the summation of the fixation probability of a BSC mutant and the fixation probability of a CeSC mutant within the entire niche, we may conclude that the optimal structure of the SC niche, which minimize the spread of both CeSC and BSC mutants in the entire niche, should have a very small number of proliferations in the CeSC compartment, i.e. 0 < *γ* ≈ 0. Furthermore, from this figure and the results provided above, we may conclude that the optimal stem cell architecture of the niche, which delays and minimizes the spread of mutants, includes a small non-zero number of migrations from BSCs to CeSCs.

When stem cells divide fully asymmetrically (*σ* = 0), then no mutants will have a chance to spread in the SC niche, i.e. the probability of fixation is zero. Simulations and analytic calculations show that when *σ* > 0, the fixation probability of mutants in the niche is not sensitive to the probability of symmetric division *σ.* However, the time to fixation is a decreasing function of *σ.* The maximum time of fixation happens when *σ* is close to zero. This result is valid for both CeSC mutants and BSC mutants. Thus, the optimal value for *σ,* which delays and lowers the spread of mutants, is zero.

## Discussion

Shahriyari and Komarova [35] have developed a bi-compartmental model for the SC niche to investigate the probability of double-hit mutant production in the niche. They found that the existence of two stem cell groups delays the double-hit mutant production. The current work generalizes that model by adding the possibility of migration from BSCs to CeSCs to study the dynamics of mutants in the SC niche. Furthermore, this work provides a framework to obtain the fixation probability of mutants in a bi-compartmental stochastic model. Analytic formulas have been obtained to calculate the fixation probability of mutants in each of compartments as well as the entire system.

The present study reveals that the small probability of migration from *S*_*b*_ to *S*_*c*_, which is reported in recent experimental observations [16], causes more delay in the process of tumorigenesis by delaying the fixation of mutants in the niche. The analytic calculations and numerical simulations, which are in perfect agreement, show that the probability of BSC mutants taking over the entire SC niche is increased when the rate of migrations from BSCs to central stem cells is increased. Additionally, the probability of CeSC mutants taking over the entire niche is decreasing when the probability of migration from BSCs to CeSCs, which is denoted by *α,* is increasing. Moreover, the time that CeSC mutants need to take over the niche is increasing when *α* is increasing. Note, the probability of mutants’ occurrences is higher in the BSC compartment comparing to the CeSC compartment [38]. Moreover, the probability that a BSC cell spreads over the SC niche is much smaller than the probability of the progeny of a CeSC mutant taking over the SC niche. Thus, we may conclude that the optimal value for *α* should be a small non-zero number.

Furthermore, the optimal structure of the SC niche must delay the spread of mutants as well as their production. It has been shown that the symmetric division of stem cells delays the production of mutants [5, 35, 37]. Moreover, the model introduced in [38] indicates that the type of stem cell divisions (symmetric and asymmetric) is not as important as the rate of divisions at each location in the tissue in terms of delaying mutants production. That model also reveals that the natural division rate of cells in the colon and intestinal crypts delays the production of mutants in the CeSC compartment. This present study shows that the location of mutants is very important in the probability that their progeny will take over the entire crypt. Mutants in the CeSC compartment have a higher chance to take over the entire crypt. Therefore, one may conclude that the natural architecture of the SC niche is the optimal structure in terms of minimizing the probability of formation of tumors.

## Materials and Methods

### Transition Probabilities

In order to calculate the fixation probabilities, we obtain all non-zero transition probabilities, which are the probabilities of moving from the state (*e*\*, *b*\*), which includes *e*\* number of *S*_*c*_ mutant and *b\** number of *S*_*b*_ mutants, to another state in an updating time step:

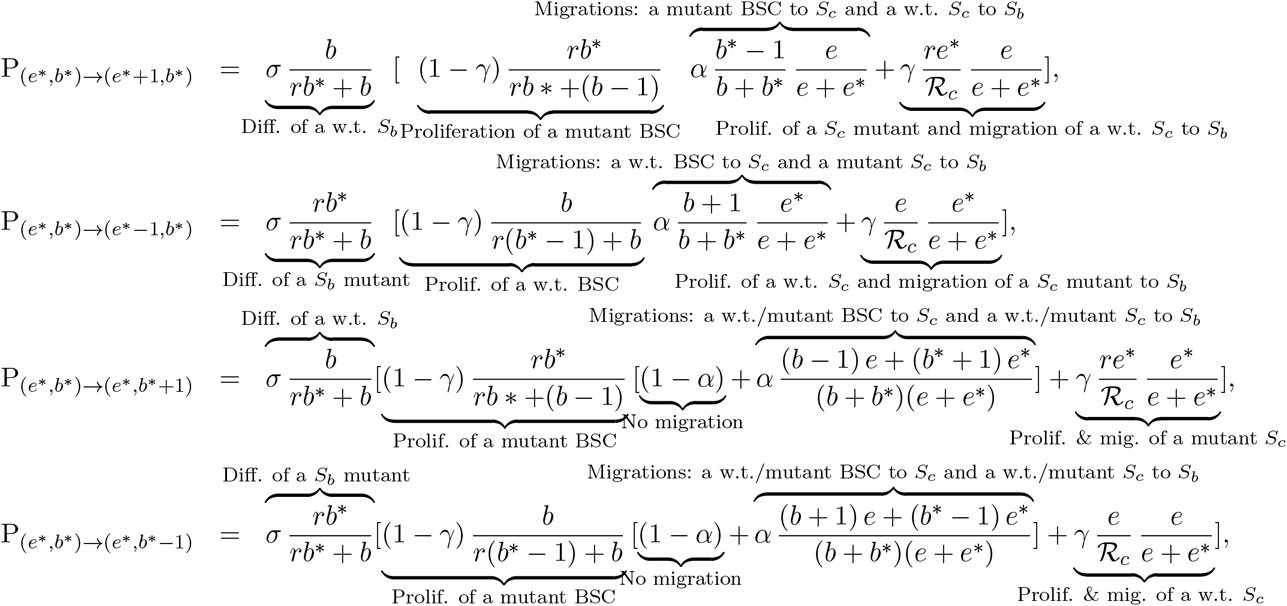

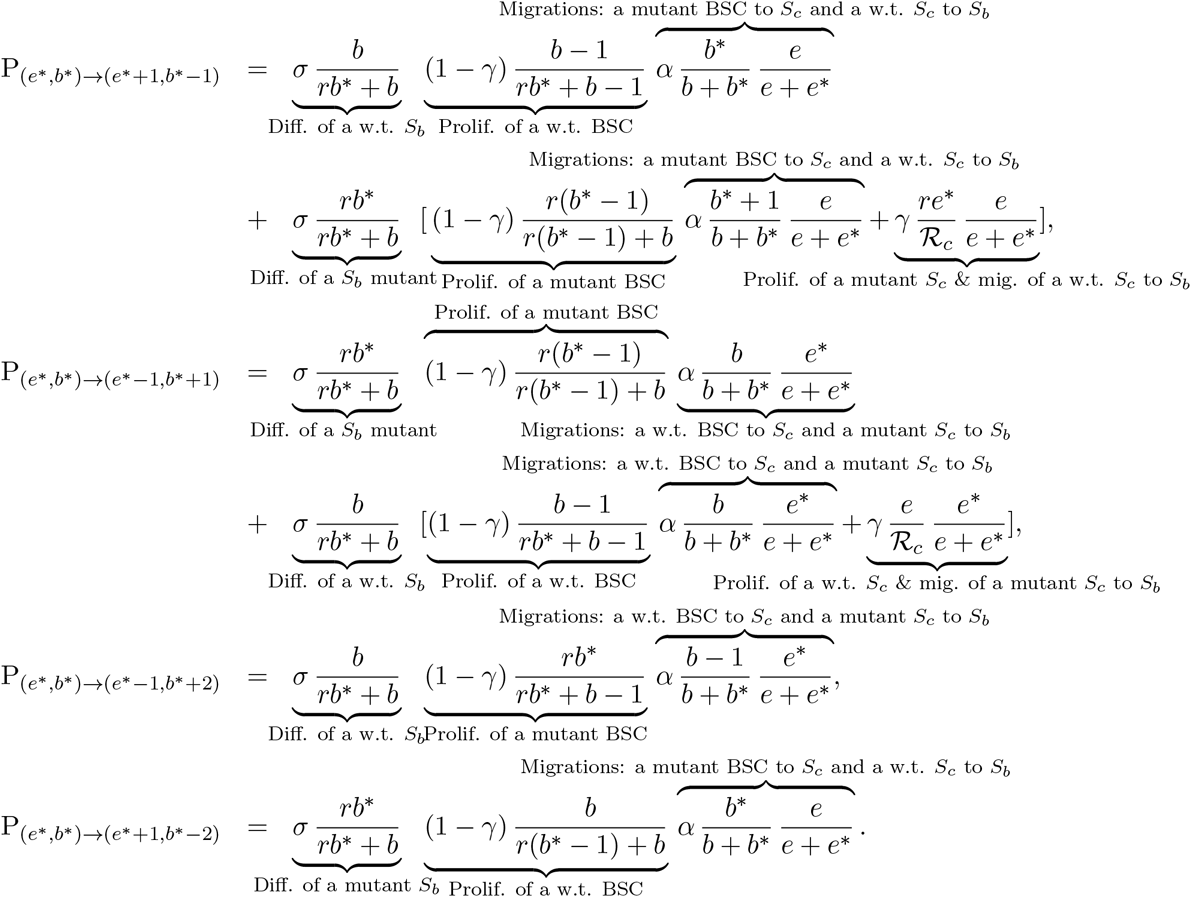

### Fixation probability of mutants in the entire SC niche

We denote the probability of the progeny of *e*\* number of *S*_*c*_ mutants and b* number of *S*_*b*_ mutants taking over the entire SC niche by π^*S*^ (*e*\*, *b*\*). The fixation probability π^*S*^ (*e*\*, *b*\*), which is the probability of moving from the state (*e*\*, *b*\*) to the state (|*S*_*c*_|, |*S*_*b*_|), i.e. all cells become mutants, satisfies the following system of equations.

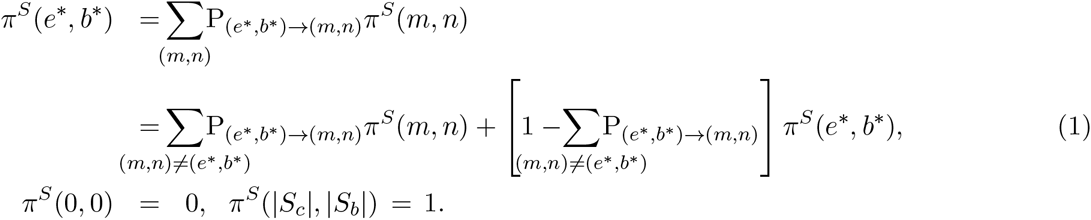

We can rewrite the above system of equations (1) in the form of

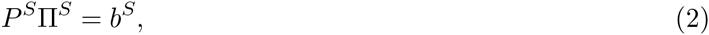

where *P*^*S*^, Π^*S*^, and *b*^*S*^ are ((|*S*_*c*_| + 1)(|*S*_*b*_| + 1) − 1) × ((|*S*_*c*_| + 1)(|*S*_*b*_| + 1) − 1), ((|*S*_*c*_| + 1)(|*S*_*b*_ + 1) − 1) × 1, and ((|*S*_*c*_| + 1)(|*S*_*b*_| + 1) − 1) × 1 dimensional matrices. We eliminate the state (0,0), because π^*S*^ (0,0) = 0. The transition probability P_(*e*\*,*b*\*)→(*m*,*n*)_ is located in the *e*\*(|*S*_*b*_| + 1) + *b*\* row and *m*(|*S*_*b*_| + 1) + *n* column of the matrix *P* if (*e*\*, *b\**) ≠ (m, n). If (*e*\*, *b*\*) = (*m, n*), the element in the *e\**(|*S*_*b*_| + 1) + *b*\* row and *e\**(|*S*_*b*_| + 1) + *b\** column of the matrix 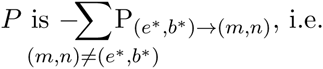.

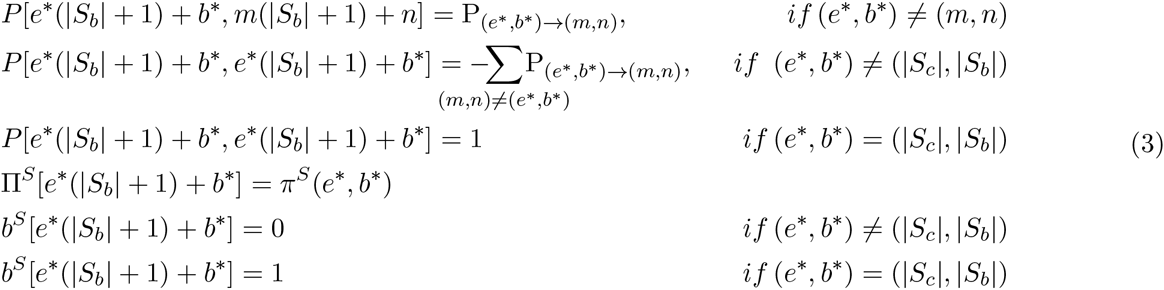

In other words,

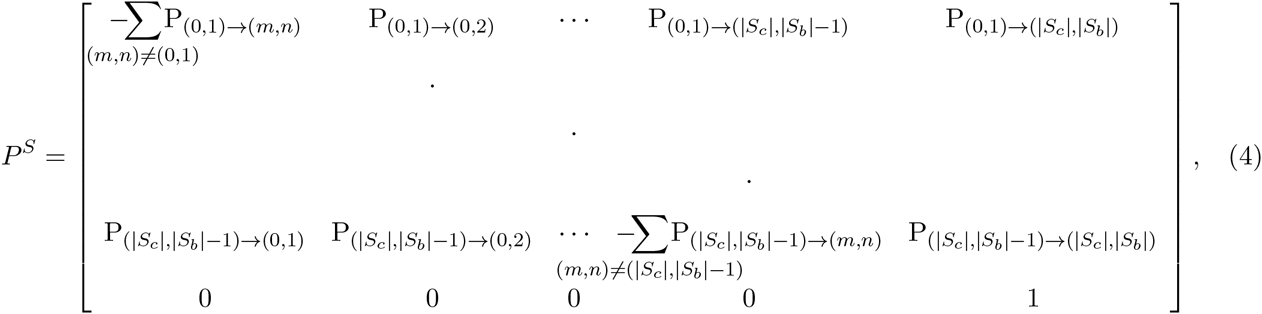

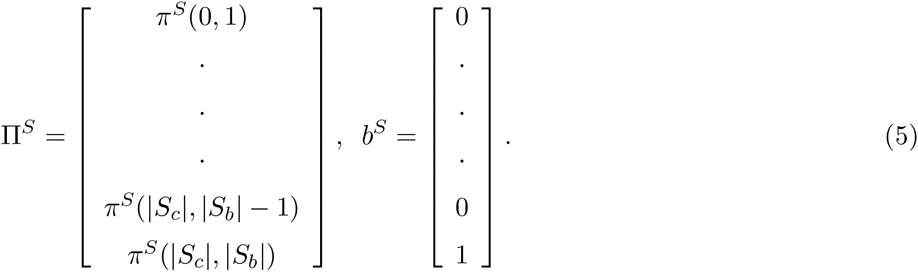

### Fixation probability of mutants in the *S*_*c*_ compartment

Following the same technique, we calculate the fixation probability in the CeSC compartment and denote the fixation probability of *e*\* number of CeSc mutants and *b\** number of BSC mutants in the CeSC compartment by π^*S_*c*_*^(*e*,b\**). The π^*S_*c*_*^(*e*,b\**) satisfies the following equations.

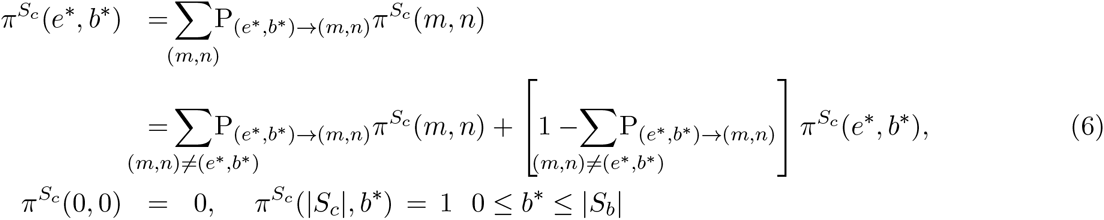

In order to find the matrix representation of the equations (6), we combine all states (|*S*_*c*_|, *b\**) 0 ≤ *b\** ≤ |*S*_*b*_| to one state. Therefore, the equations can be rewritten in the form of

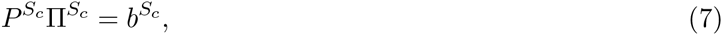

where *P*^*S*_*c*_^, Π^*S*_*c*_^, and *b*^*S*_*c*_^ are (|*S*_*c*_|(|*S*_*b*_| + 1) − 1) × (|*S*_*c*_|(|*S*_*b*_| + 1) − 1), (|*S*_*c*_|(|*S*_*b*_| + 1) − 1) × 1, and (|*S*_*c*_|(|*S*_*b*_| + 1) − 1) × 1 dimensional matrices, and

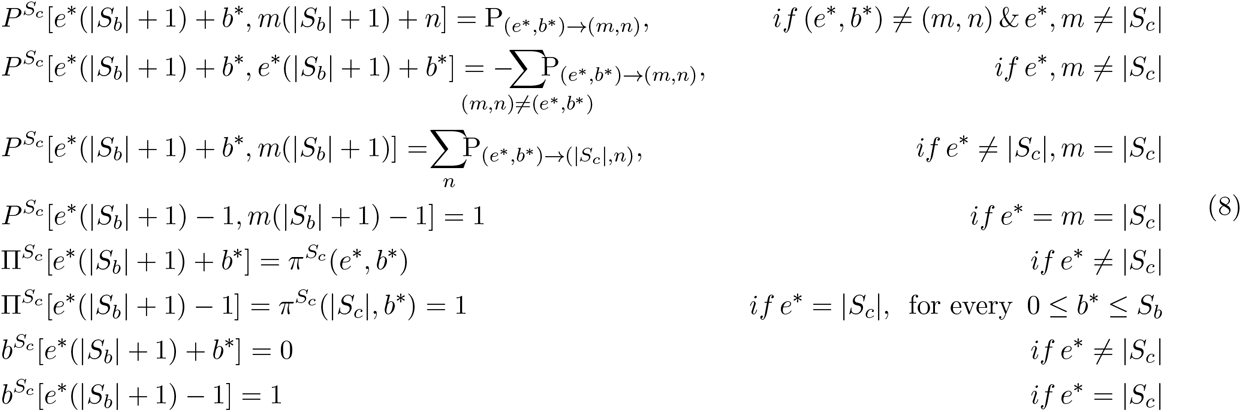

Note, the elements in the locations |*S*_*c*_|(|*S*_*b*_| + 1) + *b\** for 0 ≤ *b* ≤ S*_*b*_ in the *P*^*S*^, π^*S*^, and *b*^*S*^ matrices have been summed as one entry in the *P*^*S_*c*_*^,*π*^*S_*c*_*^, and *b*^*S_*c*_*^ matrices.

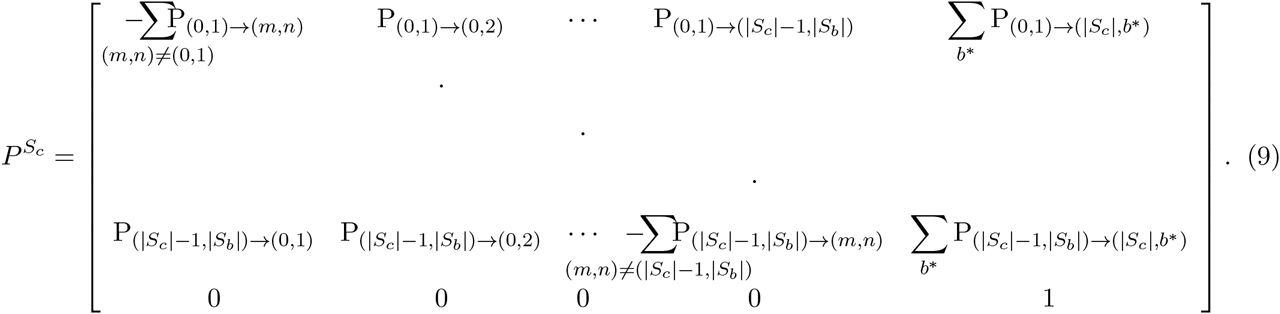

### Fixation probability in the *S*_*b*_ compartment

Here, we obtain the fixation probability, π^*S_*b*_*^(*e*,b\**) of *e*\* number of CeSC mutants and *b\** number of BSC mutants in the BSC compartment. To calculate the fixation probability π^*S_*b*_*^(*e*,b\**), which satisfies the following equation, in the BSC compartment, we combine all (*e*\*, |*S*_*c*_|) 0 ≤ *e*\* ≤ |*S*_*c*_| to one state.

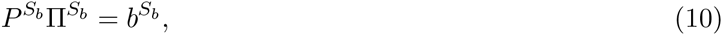

where *P*^*S_*b*_*^, *Π*^*S_*b*_*^, and *b*^*S_*b*_*^ are ((|*S*_*c*_| + 1)|*S*_*b*_| − 1) × ((|*S*_*c*_| + 1)|*S*_*b*_| − 1), ((|*S*_*c*_| + 1)|*S*_*b*_| − 1) × 1, and ((|*S*_*c*_| + 1)|*S*_*b*_| − 1) × 1 dimensional matrices, and

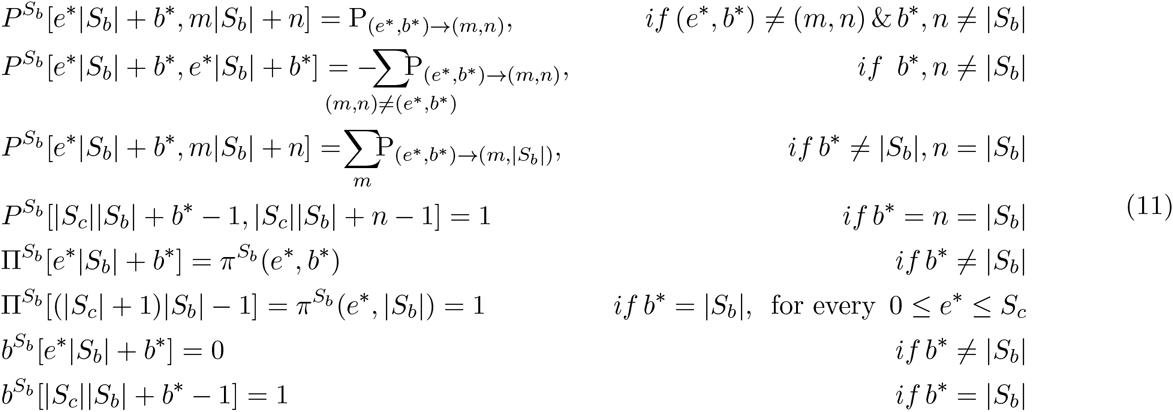

Note, the elements in the locations *e*\*|*S*_*b*_| + |*S*_*b*_| for 0 ≤ *e*\* ≤ *S*_*c*_ in the *P*^*S*^,*π*^*S*^, and *b*^*S*^ matrices have been summed up as one entry in the *P*^*S_*b*_*^,π^*S_*b*_*^, and *b*^*S_*b*_*^ matrices.

We use the matrix representation of the fixation probability equations to obtain the fixation probability of mutants at each of stem cell compartments *S*_*c*_ and *S*_*b*_ as well as the entire SC niche.

### Approximated formula

To gain more insight about the fixation probability of mutants in the entire niche, we do more analytic analysis on the equation (1). For simplicity, we assume *ℛ*_*c*_ ≈ |*S*_*c*_| and *ℛ*_*b*_ ≈ |*S*_*b*_|, and 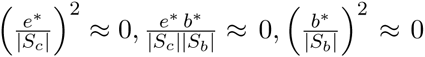. Under such a restriction of keeping the number of mutants small compared with |*S*_*c*_| and |*S*_*b*_| we can rewrite the transition probabilities as following

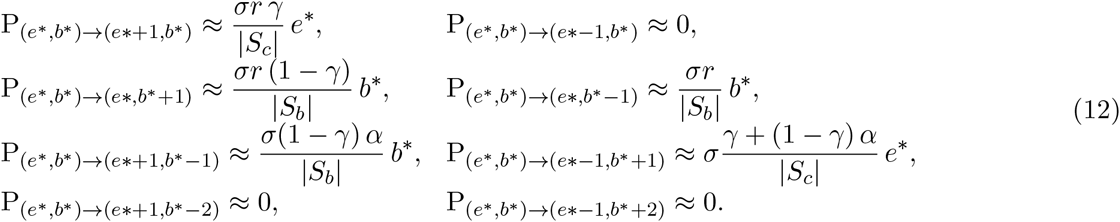

To obtain the fixation probability of mutants in the entire SC niche, we define the following generating function:

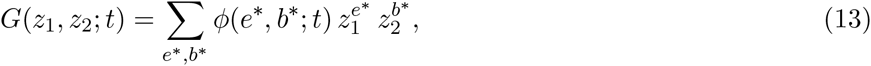

where *φ*(*e*,b*;t*) is the probability of being at the state (*e*\*,*b*\*) at time *t,* and it satisfies the following equation.

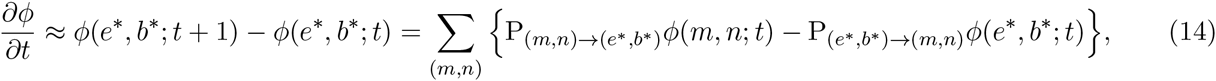

From the equations (13) and (14), the transition probabilities (12), and using the substitutions 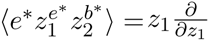 and 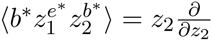, where ⟨.⟩ is the summation over states (*e*\*, *b\**) (see [39] for more details), we get

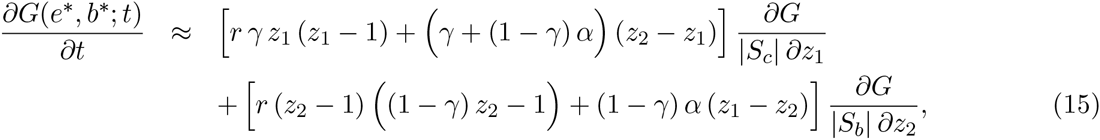

At the steady state, we can derive the following system of equations, which can provide the fixed points (in fact, the pseudo-fixed points of the system) 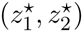 of the generating function:

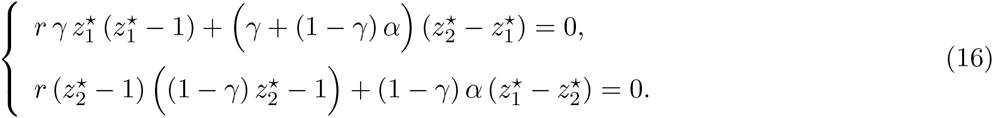

The latter system has two different solutions. The trivial solution is 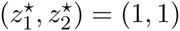, and the second solution 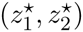 satisfies in the following equations (for *α*≠ 0,*γ* < 1):

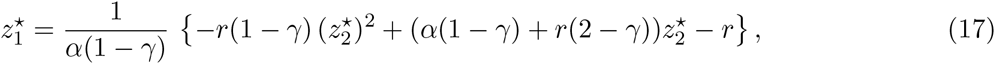

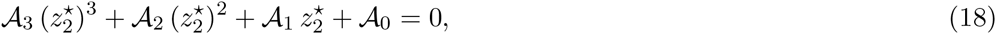

where

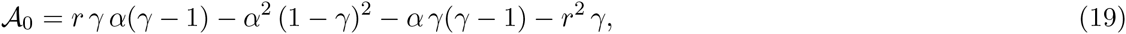

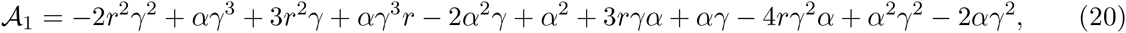

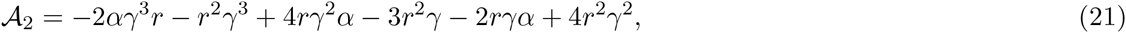

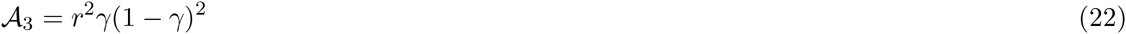

Note that when *α* = 0 then the equation (16) results in the non-trivial solutions 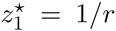 and 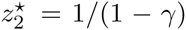. It can be easily checked that the boundary and initial conditions for the derived generating function respectively are as follows

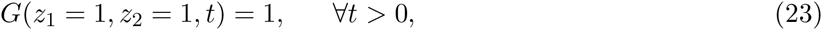

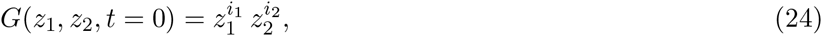

where *i*_1_ and *i*_2_ are assumed as the initial number of mutant CeSC(s) and BSC(s) respectively. The system has two distinct absorption states: the progeny of the initial mutant either fixated or eliminated from the SC population. More precisely at the steady state, we have

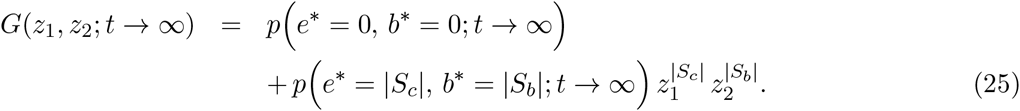

Therefore, the fixation probability of *e*\* number of CeSC mutants and *b*\* number of BSC mutants in the entire SC niche is approximately

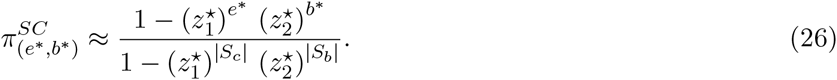

## Author Contributions

L.S. designed research; L.S. and A.M.S. performed research; L.S. and A.M.S. contributed analytic tools; L.S. performed numerical simulations; L.S. and A.M.S. analyzed data; L.S. and A.M.S. wrote the manuscript.

## Acknowledgment

This research has been supported in part by the Mathematical Biosciences Institute and the National Science Foundation under grant DMS 1440386. Authors declare that they have no competing interests.

